# A mouse bio-electronic nose for sensitive and versatile chemical detection

**DOI:** 10.1101/2020.05.06.079772

**Authors:** Erez Shor, Pedro Herrero-Vidal, Adam Dewan, Ilke Uguz, Vincenzo F. Curto, George G. Malliaras, Cristina Savin, Thomas Bozza, Dmitry Rinberg

## Abstract

When it comes to simultaneous versatility, speed, and specificity in detecting volatile chemicals, biological olfactory systems far outperform all artificial chemical detection devices. Consequently, the use of trained animals for chemical detection in security, defense, healthcare, agriculture, and other applications has grown astronomically. However, the use of animals in this capacity requires extensive training and behavior-based communication. Here we propose an alternative strategy, a bio-electronic nose, that capitalizes on the superior capability of the mammalian olfactory system, but bypasses behavioral output by reading olfactory information directly from the brain. We engineered a brain-machine interface that captures neuronal signals from an early stage of olfactory processing in awake mice, and used machine learning techniques to form a sensitive and selective chemical detector. We chronically implanted a grid electrode array on the surface of the mouse olfactory bulb and systematically recorded responses to a large battery of odorants and odorant mixtures across a wide range of concentrations. The bio-electronic nose has a comparable sensitivity to the trained animal and can detect odors on a variable background. We also introduce a novel genetic engineering approach designed to improve the sensitivity of our bio-electronic nose for specific chemical targets. Our bio-electronic nose outperforms current detection methods and unlocks a wide spectrum of civil, medical and environmental applications.

## INTRODUCTION

In the last few decades significant effort has been dedicated to developing artificial detectors for volatile organic components. Most of them are based on mass spectroscopy[1] and nano-technology[2]. However, the best chemical detectors are still the ones that emerged through biological evolution. An animal nose outperforms a majority of artificial detectors in terms of simultaneous versatility, speed, and sensitivity to specific volatile chemicals. In fact, there is greater demand than ever for the use of animals in chemical detection applications such as homeland security, defense[3], healthcare[4,5], agriculture and other fields of human activities[1]. Since the first (to the best of our knowledge) systematic training of dogs for human tracking purposes in 1899[6], animals have been employed to locate a wide variety of chemical signatures including explosives[7], illegal substances[8], bed bugs[9], and electronics[10], and to diagnose diseases such as tuberculosis[11], cancer[5,12] and Parkinson’s[13]. Despite the challenges and expenses associated with training[14], chemical detection by animals remains the gold standard in the field.

Multiple factors are responsible for the superior performance of mammalian noses in chemical detection tasks. First, biological olfactory systems are sensitive to an enormous range of structurally diverse chemicals. Over evolution, mammalian olfactory systems developed a large repertoire of olfactory receptor (OR) types (~1200 in mice), each encoded by a distinct gene[15]. Recently, it has been shown that for a given odorant, the detection threshold is defined by the most sensitive receptor[16], while an odor identity was proposed to be defined by a few most sensitive receptor types[17]. The existence of a large number of different ORs ensures high sensitivity to a broad range of different chemicals. This simultaneous breadth and sensitivity is extremely difficult, if not impossible so far, to achieve in artificial sensors.

Second, the olfactory system maximizes signal-to-noise by integrating common input from a large number of functionally identical sensors[18,19]. Each olfactory sensory neuron (OSN) in the nose expresses only one OR gene, and the axons of OSNs that express the same gene converge onto structures called glomeruli, which are arranged on the surface of the olfactory bulb—the first area that processes olfactory information. Hence, responses from a few thousands OSNs are combined and transmitted via a small number of bulb neurons (20-30 mitral/tufted cells) to high brain areas. Glomeruli provide a convenient anatomical point from which to access integrated olfactory signals from OSNs.

Third, the geometry of the nose and complex sniffing patterns solve the non-trivial problem of fast and reliable delivery and removal of odorants to and from the receptors while keeping the OSNs well-protected from the environment. This allows several important features of odor processing: fast response times on the order of ~100 ms[17,20,21], selective delivery of molecules with various chemical properties to different subsets of olfactory receptors[22,23] (but see [24,25]), and efficient odor removal from epithelium [26] between sampling events which is essential for making odor-based decisions.

Fourth, the olfactory system exhibits adaptation to background odors, facilitating robust detection of odorants in the presence of a constant or slowly varying background [27, 28], a critical function of a chemical detector.

One way to employ a biological nose for chemical detection is to use trained animals. However, training is arduous and expensive, and is usually limited to a binary reporting of the presence of only one chemical or group of chemicals [3]. Alternatively, one could potentially bypass behavior by directly recording electrophysiological responses from the intact olfactory system. Such a bio-electronic nose (BEN) would retain the benefits of the biological system, but circumvent the difficulties in measuring chemical detection behaviorally. However, reliably interfacing electronics with the olfactory system in freely moving animals and interpreting the resulting signals represent significant engineering challenges.

Here we report the development of a novel BEN based on multi-site electrophysiological recording from mouse olfactory bulb. This approach solves multiple engineering challenges, while taking advantage of the structure of the system. First, we intercept the olfactory information at the level of glomeruli, where the signals from multiple OSNs are integrated, and where activity is least affected by an animal variable behavior or internal state. Second, we utilize surface grid electrodes to detect odor-evoked local field potentials[29], which are stable over many months of recording—something that is difficult to achieve with depth electrodes. In principle, one could use glomerular calcium imaging as a sensitive readout of glomerular activity[16]. However, imaging would be harder to implement in in freely moving animals in future applications. Third, have chosen the mouse as model organism for the BEN in order to capitalize on genetic methods that can be used to modify the repertoire of olfactory receptor genes and the arrangement of glomeruli to tune the system for specific tasks.

Our BEN was able to detect chemicals at concentrations comparable with animal detection thresholds. We tested the system for odor detection in the presence of background odor, and explored the possibility of using genetically modified mice to enhance BEN detection capabilities for specific odors. Using mice that overexpress a specific receptor type improved detection in the presence of masking odors, without affecting overall sensitivity. This series of experiments lays the foundation for using BEN systems in a variety of applications including advanced chemical detection.

## RESULTS

### Basic characterization of the odor related signals

To measure odor-driven spatiotemporal patterns of neuronal activity from the intact olfactory system, we chronically implanted mice with a 64-site surface electrode array positioned on the dorsal surface of the olfactory bulb (Fig. 1A). We recorded local field potentials (LFP) from the surface of the bulb in awake, head-fixed mice while presenting to the animals a series of monomolecular odorants and odorant mixtures at different concentrations. To monitor the actual delivery of odors to the nasal cavity, we recorded sniffing patterns via an external pressure sensor located in the odor port. Each functional electrode exhibited a common signal as well as site-specific components (Fig. 1A). Both signals carried information about odor identity (Fig 1B) and odor concentration (Fig. 1C), with the common signal stable over many weeks of recordings (Fig 1B).

**Figure 1.**
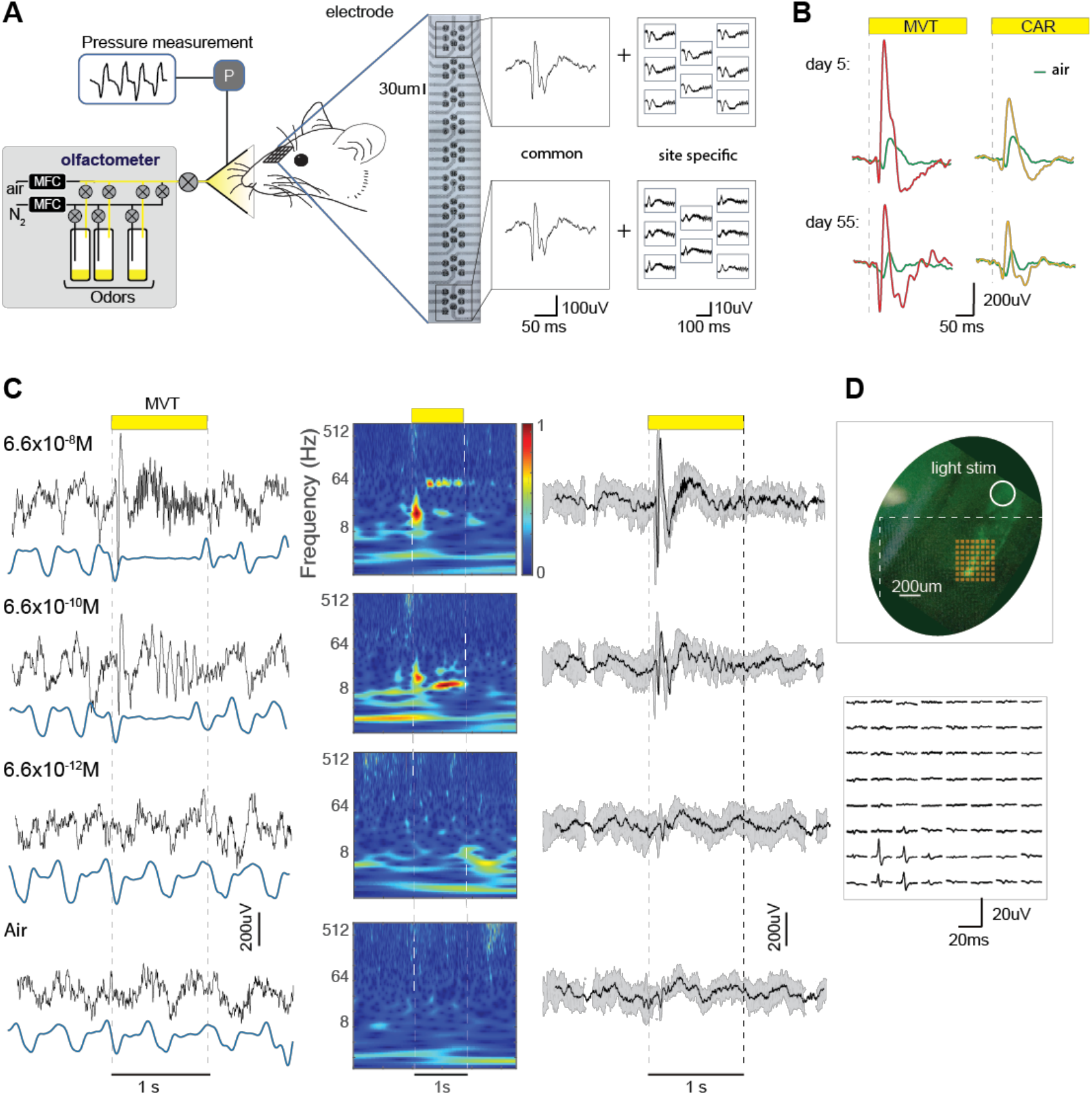
Bio-electronic nose design and input signal characterization. **A.** System design. A mouse is exposed to various odors at different concentrations delivered from an odor port using an air-dilution olfactometer that regulates flow rates with mass flow controllers (MFC). The sniff pattern is monitored by pressure sensor (P) in the odor port. A grid electrode with 64 sites is chronically implanted on the mouse olfactory bulb. The LFP signals from the electrodes can be decomposed into a common and site-specific component. **B.** Example common signals for two different odors methyl valerate (MVT), and carvone (CAR), recorded 5 and 55 days after implantation. **C**. Typical LFP signal for multiple MVT concentrations and air. *Left*: single trial LFP traces (black) and sniff (blue), middle: corresponding LFP spectrograms, *right*: average LFP (n = 50 trials) aligned to inhalation onset. Odor delivery marked by yellow bars and dashed lines. **D.** Optogenetic stimulation of a single glomerulus. *Top*: image of the olfactory bulb with a single glomerulus expressing ChR2 and YFP (see Methods), with a grid electrode array positioned on the surface of the bulb. The dashed lines indicate the edges of the device, and small squares are individual electrode sites. The circle indicates the position of the light spot for illumination of the OSNs axons converging on the photoactivatable glomerulus. *Bottom*: site specific signals in response to light stimulation.

Odor stimuli usually evoked a transient LFP signal with a typical duration of ~150 msec from the onset of the first inhalation after odor delivery. This transient response was followed by LFP oscillations, in a frequency range which depended on odor concentration, as previously reported [30]: high concentrations usually evoked a γ-range oscillation (~60Hz), and lower concentrations evoked β-range oscillation (~8Hz). Θ oscillations (~3Hz) were present throughout the recordings (Fig. 1C).

To demonstrate the ability of individual electrodes to detect activity from individual glomeruli, we selectively stimulated a single glomerulus using optogenetics[31]. We used a strain of mice (M72S50-ChR2) in which a single population of OSNs (and thus a set of single glomeruli) can be activated with light based on expression of channelrhodopsin. We shined light on the axons of these OSNs and recorded signals from an electrode array which covered the target glomerulus. We observed that only the few electrode sites that were in close proximity to the activated glomerulus recorded a light triggered LFP signal (Fig. 1E). These data show that activation of individual glomeruli evoke spatially localized LFP signal.

### The bio-electronic nose accurately classifies odor identities

To determine to which degree LFP signals from the olfactory bulb could be used to identify different chemicals, we stimulated the BEN with a set of individual odors—methyl valerate (MVT), ethyl tiglate (ETG), hexanal (HEX), carvone (CAR), and benzaldehyde (BZD). The odorant set included chemicals form the same (ETG and MVT) and different groups (CAR and MVT), and those which activate predominantly dorsal (MVT and BZD) and ventral (CAR) glomeruli.

For the proof-of-principle approach, we first performed classification analysis based only on the signal during the first inhalation after odor onset, by selecting a time window of 300 ms aligned to the first inhalation onset and discarding data from subsequent sniff cycles. To identify a robust feature space for classification we compared different representations of the data. The cross-electrode, common signal, could classify 6 stimuli (5 odorants and air) with an average accuracy 71±4% (n = 6 mice, mean ±STD, cross-validated; see Methods). A classifier based on the site-specific signals achieved 82±12% accuracy. Combining two sets of features further improved the performance to 83±11%.

The modest improvement in performance of the classifier that uses the site-specific signal compared to one that only uses the common signal may be owed to a combination of the small amount of data paired with a significant increase in model complexity (number of parameters). To overcome this limitation, we used dimensionality reduction techniques to project the site-specific signals into a 5-dimensional latent space, explaining an average 87% of the signal variance. Using this representation, we were able to achieve the similarly high performance, 83±8% (Fig. 2B and Suppl. Fig. 1A), while substantially reducing data and computational requirements.

**Figure 2:**
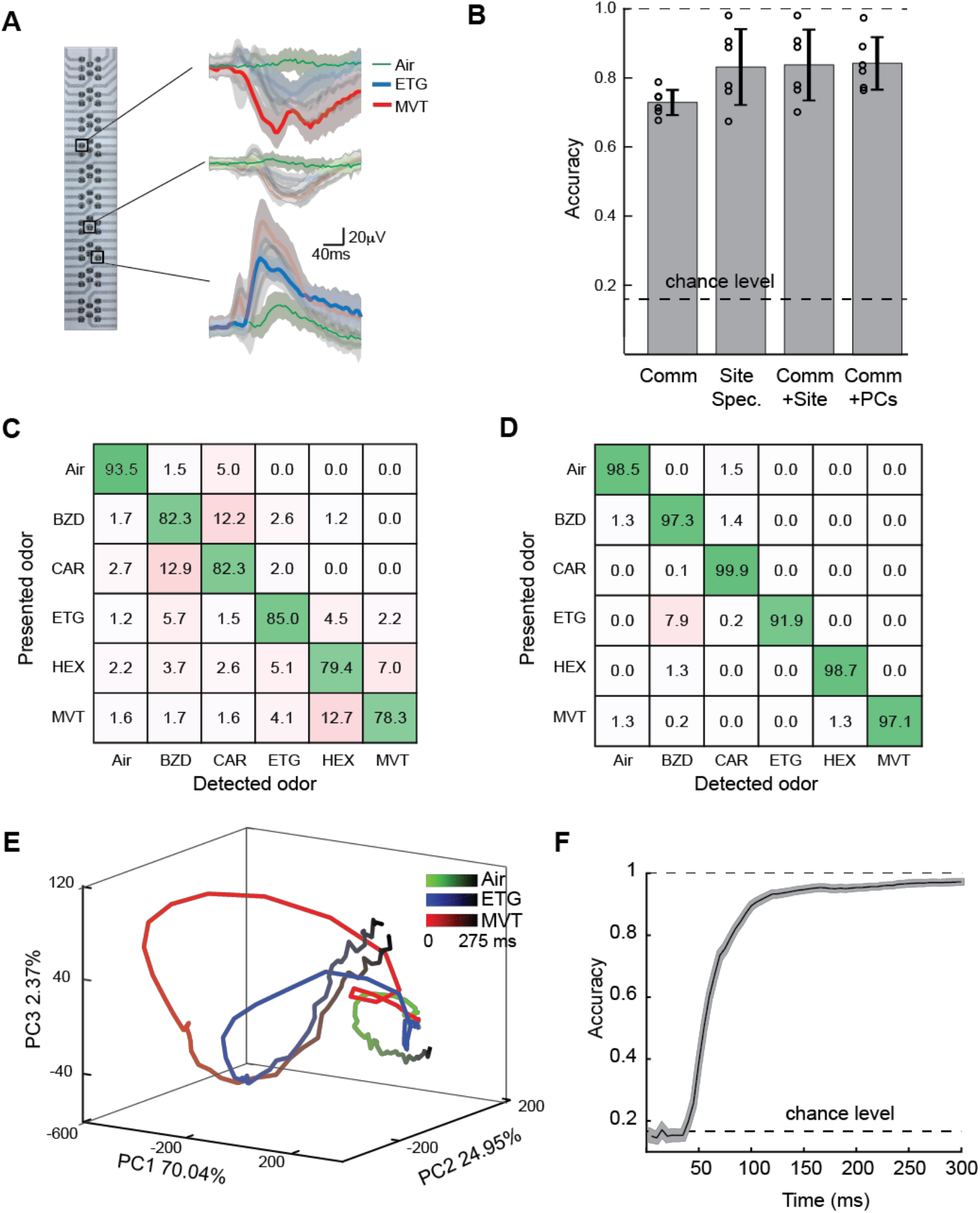
Bio-electronic nose odor identity classification. **A.** Site specific responses from representative electrodes for ethyl tiglate (ETG, blue), methyl valerate (MVT, red), and air (green). Shaded areas indicate standard deviation (s.d.) across trails. **B**. Classification performance for 6 stimuli (5 odors and air) using only common signal (Comm), only site-specific signals (Site Spec.), both (Comm+Site), and common signals and the first 5 PCs of site-specific signals (Comm+PC). Circles correspond to individual mice (n=6); error bars show s.d. **C, D.** An average (**C**) and the best-performer (**D**) confusion matrices for n=6 mice presented with a battery of 6 different stimuli (classified using Comm+PCs). **E**. Temporal trajectories of site-specific signals in the first 3 PC dimensions, for different odorant. Later time along trajectories is encoded by darker color. **F.** Classification performance for increasing time windows from stimulus onset. Shaded area indicates 95% confidence bounds, bootstrapping averaged over 100 samples.

The confusion matrix provides additional information about the nature of the errors of the classifier (averaged across all animals, Fig. 2C, and the best performing animal, Fig. 2D). The matrix provides information about which errors were most prominent for specific odor classifications. Even for a ventral odor, such as CAR, the overall classification accuracy was 82.3%, and its highest misclassification error was with BZD, which is a dorsal odor. Odors for the same chemical group (ETG and MVT) were rarely misclassified (2.2% and 4.1%). Dorsal odors from different groups (CAR and MVT) were also rarely misclassified (1.6% and 0.0%), suggesting good coverage of activated glomeruli improve the recorded signal to noise ratio.

To characterize the temporal extent of odor specific information, we visualized the response of the 64 electrodes to specific odors as trajectories in the space of the first three PCs. The trajectories for individual odors start at the same fixed point and diverge after ~50 ms (Fig. 2E) and remained distinguishable for the subsequent 200 ms. Reflecting this temporal structure, decoding odor identity in increasingly large windows starting from the onset of inhalation reveals a sharp transition from chance levels to asymptotic decoding performance in the window 50-100ms (Fig. 2F). This time course was similar in the entire cohort of mice recorded (Suppl. Fig. 1B). The predictive power of the early version of the signal justifies our choice of focusing on the first inhalation and ignoring slower oscillatory components in the signal (Fig. 1C).

### Detection accuracy matches behavioral thresholds

To investigate how concentration affects odor detection, we recorded neural signals in response to single odorants presented across a wide range of concentrations. We observed that changes in concentration resulted in a qualitative scaling of the signal, with higher concentrations giving higher the signal amplitude and shorter response latency (Figs.1C and 3A). As a concentration is a continuous variable, we used linear regression to predict the concentration from the LFP signal (Fig. 3B, see methods). The average concentration error was around 1 order of magnitude (1.1 in log10 units) for most of the concentrations. The precision was lost at very low concentrations of ~10^−16^. The results for classification analysis for multiple concentrations similar to that for multiple odorants are in Sup. Fig. 2.

**Figure 3:**
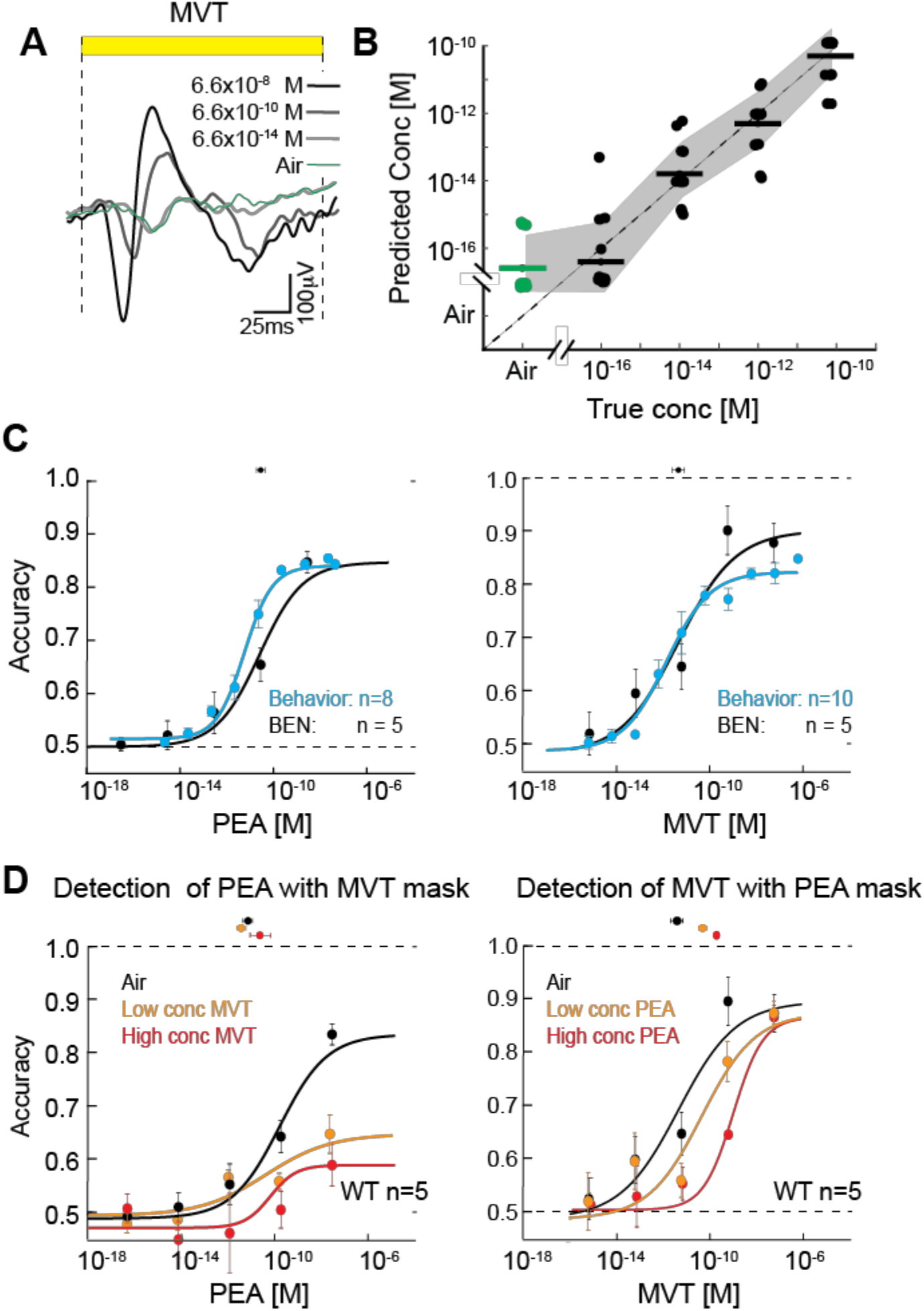
Classification accuracy and sensitivity for different concentrations. **A.** Average signal of a representative electrode site for different concentrations of MVT and air (n=20 trials per concentration). **B.** Comparison of BEN concentration estimates (linear regression) to true concentration, horizontal bars and dots indicate averaged and individual animal predictions, shaded area correspond to ±1 s.d. **C.** Average accuracy for mouse behavioral performance in odor detection task as a function of odor concentration (blue) for two odors PEA (left) and MVT (right), and BEN performance in the same conditions (black). Lines are model fits (see Methods). **D**. BEN average odor detection accuracy in the presence of a masking odor (n = 5 mice) for detection of PEA in the presence of MVT (left) and detection of MVT in the presence of PEA (right), without a mask (black), and for low (orange) and high (red) mask concentrations. Maximum performance for behavioral thresholding is 85% [16]. EC50 values are indicated above each plot in C and D.

Next, we compared the sensitivity of the BEN to that of intact, behaving mice using two odorants: MVT, an ester that generates broad patterns of activity in the olfactory bulb, and PEA, an amine that specifically activates a small subset of dorsal glomeruli [16]. An important feature of choosing PEA is that the glomerulus corresponding to the most sensitive receptor TAAR4, is positioned on the dorsal surface of the bulb and accessible to the electrode [16, 32]. Moreover, behavioral detection thresholds have been carefully measured for both MVT and PEA using a go/no-go thresholding paradigm [16]. The average behavioral detection accuracies (PEA: n = 8, and MVT: n = 10) from this previous work are shown on Fig. 3C (blue dots and lines). At the highest concentrations, mice reach above 80% accuracy in detecting a target odor. The maximum performance in the task is 85%; see [16], and the sensitivity is defined as the concentration at half-maximal performance (EC_50_beh_). These were EC_50_beh_ = 5.0 × 10^−12^ M for PEA and EC_50_beh_ = 1.7 × 10^−12^ M for MVT [16].

To define the relative sensitivity of the BEN, we trained the classifier to discriminate between an odorant at any non-zero concentration versus air, using the same data as for concentration classification analysis for PEA and MVT. For both odorants the classification accuracy was high for high concentrations and dropped to chance as the presented concentration dropped. For MVT, the sensitivity of the BEN (EC_50_BEN_ = 4.8 ± 0.3 ×10^−12^ M, mean±SD) matched that for the behavioral performance (sensitivity = 1.7 ×10^−12^ M, [16]). For PEA, the sensitivity (EC_50_BEN_ = 3.1 ± 0.2 × 10^−11^ M) was within one order of magnitude of the behavioral threshold (sensitivity = 5.0 × 10^−12^ M, [16]) (Fig. 3C). Thus, the odor detection performance of the BEN closely matches that of mouse behavior.

### Detection accuracy persists in the presence of background odor

Under natural conditions, monomolecular odors are rarely present in isolation. To test the BEN under more realistic conditions, we measured the detection accuracy of the same odorants as mixtures, to mimic a target odor masked by a background odor.

First, MVT was masked with two concentrations of PEA: a low concentration mask of 6.6 × 10^−10^ M and a high concentration mask of 6.6 × 10^−8^ M. We then compared both the MVT detection threshold and detection accuracy in the presence or in the absence of the masking odor (i.e. in air alone). We observed that as the concentration of masking odor increased, the sensitivity decreased (EC_50_= 4.8 ± 0.3 ×10^−12^ M for no mask, to 6.6 ± 0.3 ×10^−11^ M for the low concentration mask, and 7.6 ± 0.1 ×10^−11^ M for the high concentration mask). However, in all conditions, the highest detection accuracy values were comparable (accuracy = 90±9% no mask, 89±8% high PEA mask, and 89±12% low PEA mask, comparison between no mask and low/high concentration mask: *p*≈0.48/0.50, two-tailed t-test)

The results were different for detection of PEA in the presence or absence of the masking odor MVT (low concentration mask = 3.5 ×10^−12^ M; high concentration mask = 3.5 ×10^−10^ M). Here, the EC_50_ values for PEA detection were comparable among all three groups: (EC_50_ = 3.1±0.2 ×10^−11^ M for no mask, 1.3±0.4 ×10^−11^ M for low concentration mask, and 4.8±0.4 ×10^−11^ M for high concentration mask). However, the maximum accuracy of PEA detection dropped from 85±3% to 65±4% and 60±9% for low and high mask concentrations, respectively (comparison between no mask and low/high concentration mask: *p* < 0.02/0.0002, two-tailed t-test) (Fig. 3D).

One possible explanation for this difference in effect may relate to the patterns of glomerular activation elicited by the two odorants (see Discussion). Overall, the data show that the BEN can detect and classify a target odor in a mixture—a step towards real-world applications.

### Receptor overexpression improves BEN detection accuracy

One of the unique features of a mouse-based BEN is our ability to genetically modify the composition of OSNs in the olfactory epithelium, and to tune the system to specific odorants. To begin exploring this capability, we tested the effect of increasing the number of OSNs (and glomeruli) that respond to a given odor on BEN detection. To do this, we exploited the fact that the most sensitive receptor for PEA is known [16]. We hypothesized that increasing the number of glomeruli corresponding to this receptor (TAAR4) might improve the detection of PEA, while not changing the detection of MVT (to which TAAR4 does not respond). To test this prediction, we used transgenic mice that have an increased number of TAAR4-expressing OSNs, leading to a ~20-fold increase in the number of TAAR4 glomeruli ([16]; Fig. 4A). We compared the BEN performance for wild-type (WT) and TAAR4 over-expressing (OE) animals for detection of PEA and MVT, with and without masking.

**Figure 4:**
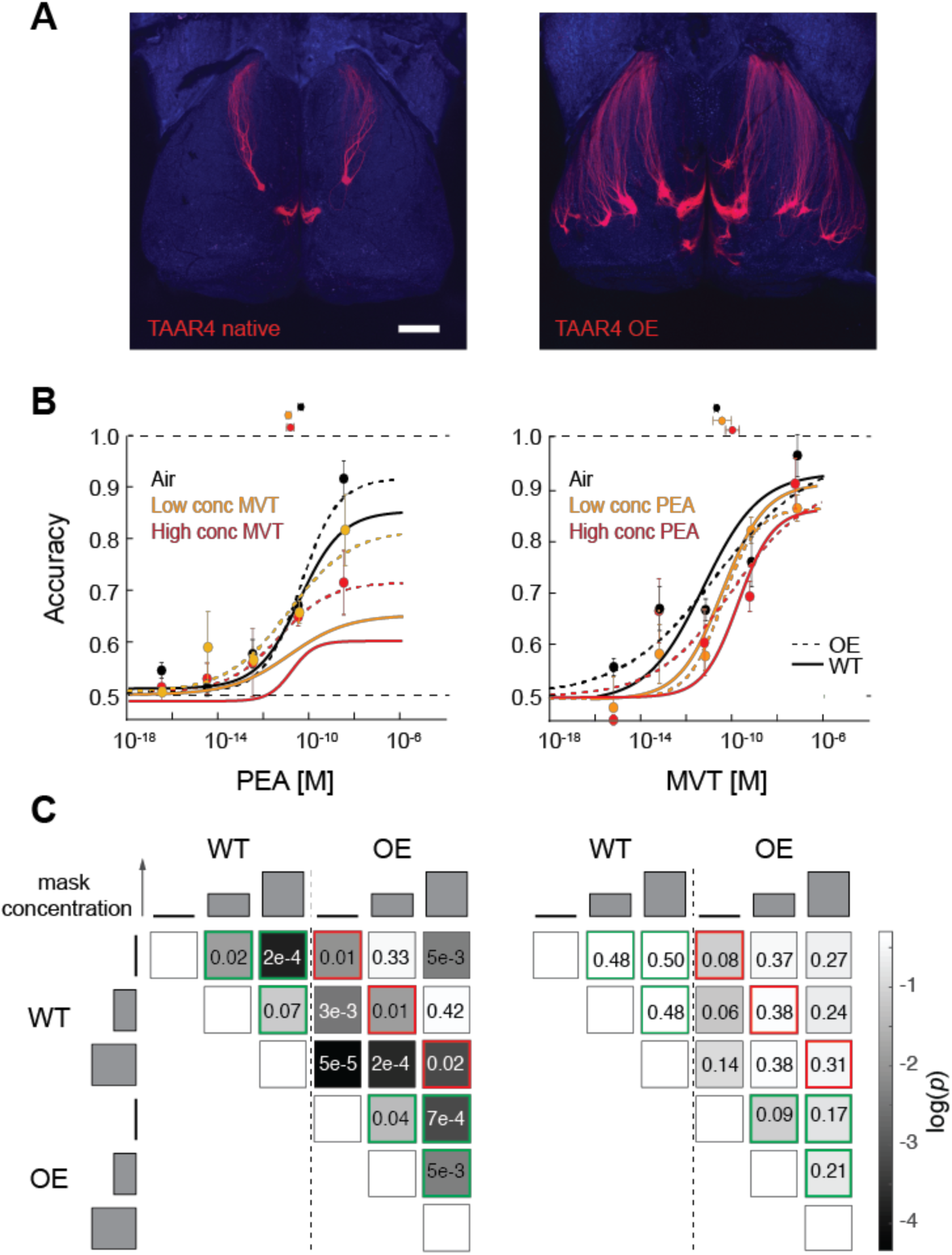
Performance of BEN using mice overexpressing TAAR4 receptors. **A**. Fluorescent image of olfactory bulbs of unmodified, native (WT, left) and overexpresser (OE, right) animals. All OSN expressing TAAR4 receptors also expressed RFP. Scale bar = 500um **B**. BEN odor detection accuracy in the presence of a masking odor for OE mice (circles - average performance for *n*=4 mice, error bars - ±1s.d, dashed lines – logistic fits) and for WT mice (solid line – logistic fits from Fig. 3D, data points are omitted, n=4) for detection of PEA in the presence of MVT (left) and detection of MVT in the presence of PEA (right), without a mask (black), and for low (orange) and high (red) mask concentrations. **C.** Summary comparison of maximum detection accuracies across various conditions using two-tailed t-test p-values for PEA detection with MVT mask and MVT detection with PEA mask: no mask (air), medium mask concentration, and high mask concentration, and for wild type (WT, n=5) and overexpressor (OE, n=4) mice (see methods). The effects of masking are shown in green contoured squares, and the effects of WT vs OE for the same masking conditions are shown in red contoured squares.

We observed that detection of PEA (both with and without masking odor) improved in OE animals compared to WT. The maximum detection accuracy for PEA with no mask increased from 85±3% for WT to 92±8% for OE mice (p< 0.01). Similarly, the maximum detection accuracy for the low concentration mask increased from 65±4% for WT to 82±16% for OE mice (p<0.01), while accuracy increased from 60±9% to 70±12% for the high concentration mask (p<0.02) (Fig. 4B left, Fig. 4C bottom). At the same time, overall sensitivity was the same: EC_50_ values were comparable at 5.4±0.1 ×10^−11^ M, 1.2±0.1 ×10^−11^ M and 1.58±0.2 ×10^−11^ M for no mask, low and high concentration masks respectively.

On the other hand, overexpression of TAAR4 did not affect the maximum detection accuracy for MVT. The maximum accuracies for WT and OE mice with no mask were 90±9% and 92±3% (p ≈ 0.08), with a low concentration mask: 89±8% to 90±2% (p ≈ 0.38), and with a high concentration mask: 89±12% to 89±7% (p ≈ 0.31).

For the same comparison, overall sensitivity did shift. The EC_50_ for MVT decreased from 4.8±0.4 ×10^−12^ M in WT to 2.0±0.1 ×10^−11^ M in OE mice in the absence of masking odor. In the presence of masking odor, the effect was weaker: EC_50_ = 6.6±0.3 ×10^−12^ M for WT and 7.6±0.1 ×10^−11^ M for OE (low PEA mask) and EC_50_ = 4.1±0.2 ×10^−11^ M for WT and 8.9±0.1 ×10^−10^ M for OE (high PEA mask).

To quantify overall detection performance across groups, we compared accuracies across conditions at the high odor concentrations within and across animal groups (Fig. 4C). We found systematic improvements in performance for the same masking condition for OE relative to WT, for detecting PEA in MVT mask, but not for detecting MVT in PEA mask (red contoured squares, Fig. 4C). In addition, for PEA but not MVT masking considerably impacts BEN performance (green contoured squares, Fig. 4C).

Thus, receptor overexpression improves overall performance of the BEN for detection of monomolecular odors and odors on a background, with odor-specific increases in sensitivity and maximum accuracy.

Overall, the detection sensitivity of the BEN for pure odorants matches behavioral thresholds. While masking odorants impact sensitivity, overexpressing a high affinity receptor significantly improves the detection of a target odor in the presence of masking odors, especially in cases where it is difficult to obtain good electrode coverage of the glomeruli that are activated by a desired target.

## DISCUSSION

Here we report an approach to exploit the mouse olfactory system for sensitive and versatile chemical detection. Our bio-electronic nose is capable of discriminating among multiple presented odorants (Fig.2), estimating odorant concentration (Fig.3), and detecting odorants with sensitivities that are comparable with those of well-trained animals (Fig.3). Moreover, the BEN performed well when faced with one of the hardest problems in chemical detection --odor identification in the presence of masking or background odors. Even in the presence of a masking odor, the BEN performance remained significantly above chance. One promising direction for improvement capitalizes on our choice of animal model, the mouse, which benefits from the availability of modern genetic tools. We demonstrated that genetic manipulation, leading to overexpression of a specific OR, improved detection of target odors in the presence of an odor background (Fig. 4).

While our implementation of the BEN requires further optimization, it has shown remarkable performance in initial tests. This performance is notable because it is unlikely that the BEN has direct access to all relevant olfactory sensory inputs. When a trained animal detects a specific odor, the brain has access to information flowing into all olfactory bulb glomeruli. In contrast, the BEN presumably records only some portion of that incoming information, perhaps restricted to the activity in the directly underlying dorsal glomeruli. It was, therefore, not obvious a priori that the performance of the BEN could rival that of the intact system. Despite this concern, the sensitivity of the BEN was similar to that of behaving animals. Prior work has shown that behavioral detection thresholds are defined by the activation of the most sensitive glomeruli[16]. In our experiments, the electrode array was positioned on the OB in a way that most likely covered the glomerulus corresponding to the most sensitive receptor for PEA, TAAR4[33]. This may explain why the BEN was able to detect PEA at such low concentrations.

For MVT, we do not know the identity or location of the most sensitive glomerulus. Even if the most sensitive MVT glomeruli were not covered directly by the array, it is possible that activation of remote glomeruli could create broader perturbations of the LFP, which might then be picked up by underlying electrodes. Such “indirect” detection of signals entering through glomeruli that are far from the array could significantly expand the spectrum of chemicals that can be detected at low concentrations. Our demonstration that the BEN can exhibit high sensitivity is critical for a large variety of applications.

Our data indicate that the earliest activated glomeruli play a particularly important role for BEN performance. We observed that discrimination accuracy for multiple odorants increased as a function of time (Fig. 2F) and saturated quickly (~100 ms), much faster than the activation time course for a majority of glomeruli[34] Fig. 4). The fastest glomeruli to respond may correspond to the most sensitive receptors, which are sufficient to reach high levels of odor discrimination[17]. Further experiments will demonstrate how performance of larger electrode arrays, which cover larger numbers of highest affinity receptors improve detection and discrimination for a large number of odorants.

We also observed that MVT detection is less affected by PEA as a masking odor, compared to PEA detection when using an MVT mask (Fig. 3D). One possible reason for this asymmetry is that MVT may excite a larger number of glomeruli than PEA at the concentrations we tested[16,32,33,35]. This explanation is also consistent with the fact that when the number of glomeruli activated by PEA is increased via genetic modification, the effect of the MVT mask is suppressed (Fig. 4B&C). These results are crucial for developing future strategies to improve detection of defined odors in the presence of different backgrounds, and underscore the importance of covering a larger number of glomeruli as a pathway towards improving BEN performance.

Our BEN approach allows us to read complex olfactory information from the intact olfactory system rapidly and reliably. This opens up the possibility of using this approach to detect specific chemical targets under more real-world conditions. A potential downside of our approach is that it requires the use of animals and implantation of a brain machine interface. This may limit its application in some situations. However, trained animals are frequently used in chemical detection[36]. Our system provides a method to use biological noses in a way that does not require exhaustive training.

More broadly, the fact we now possess a setup that translates odor responses into complex data sets may enable us to discriminate among a large array of chemical targets. For example, data from the BEN might be used to train AI algorithms to discriminate odor samples taken from hospital patients with known medical conditions, and the database used to categorize novel samples. This process could yield the odor footprints of various diseases and provide a cost effective, non-invasive broad-spectrum diagnosis method.

## MATERIALS AND METHODS

### Animals

For basic electrophysiological experiments, we used 11 adult homozygous *M72–IRES-ChR2-YFP* mice (Strain *Olfr160 tm1.1(COP4*/EYFP)Tboz, males*). For experiments with receptor overexpression we used 7 mice that overexpress TAAR4 receptors (5×21-TAAR4Tg [16]). For optogenetic stimulation of a single glomerulus we used one male M72/S50-IRES-tauGFP mouse (strain *Olfr545 tm3(Olfr160)Mom*). For behavioral experiments we used C57BL/6J mice. Animals were 6–10 weeks old at the beginning of experiment and were maintained on a 12-h light/dark cycle (lights on at 8:00 p.m.) in isolated cages in a temperature- and humidity-controlled animal facility. All animal care and experimental procedures were in strict accordance with protocols approved by the New York University Langone Medical Center and Northwestern University Institutional Animal Care and Use Committees.

### Chronic electrode implantation

Mice were anesthetized with isoflurane (2-3%) in oxygen and administered ketoprofen (0.1 mg/kg) as analgesic. The animals were secured in a stereotaxic head holder (Kopf). After incision of the scalp the connective tissue covering the skull was removed with H_2_O_2_ (5%). One micro screw was placed into the skull at 1mm caudal to lambda. A custom-built plastic 3D-printed head-bar [37] was attached to the skull using Vetbond cyanoacrylate glue. Head-bar and ground screw were cemented in place using dental cement (Dental Cement, Pearson Dental Supply). The skull was thinned and a small craniotomy was performed at the site of the electrode implantation. The surface Electrode (Diagnostic Biochips or Malliaras Lab, Cambridge, UK) then was placed on the bulb. To achieve consistency of placing electrodes across mice, we used M72 fluorescent glomerulus in M72-ChR2 mice as a landmark. For a single glomerulus stimulation experiment, using M72-S50-ChR2 mice we insure that electrode covered the fluorescent glomerulus. Following placement, the electrode was secured with Kwik Sil (World Precision Instruments). After the Kwik Sil cured the electrode, PCB was attached to the head bar using 5-min epoxy glue. The electrode surgery site was then sealed with Body double mold rubber (Smooth-On, Easton PA). After surgery, mice were individually housed and given at least two days for recovery before water deprivation or data recording.

### Odorants

The odorants used in the experiments are listed in Table 1. The odorants were diluted in water and kept in dark vials (45 mL volume filed with 5 mL diluted odorant). Dilutions for concentration series (Figs. 3 and 4) were prepared by subsequent dilutions of the freshly made PEA and MVT odorants. Prior to odor delivery, the headspace odorant was additionally diluted from 10 to 100 folds.

**Table 1.** Odorants used in the experiments

| Odorant | Abbreviation | Dilution | Headspace concentration |
| --- | --- | --- | --- |
| Phenyl-ethylamine | PEA | 2x10<sup>-2</sup> | 4.4x10<sup>-8</sup> M |
|  |  | 2x10<sup>-4</sup> | 4.4x10<sup>-10</sup> M |
|  |  | 2x10<sup>-6</sup> | 4.4x10<sup>-12</sup> M |
|  |  | 2x10<sup>-8</sup> | 4.4x10<sup>-14</sup> M |
|  |  | 2x10<sup>-10</sup> | 4.4x10<sup>-16</sup> M |
| Methyl valerate | MVT | 8x10<sup>-2</sup> | 7x10<sup>-6</sup> M |
|  |  | 8x10<sup>-4</sup> | 7x10<sup>-8</sup> M |
|  |  | 8x10<sup>-6</sup> | 7x10<sup>-10</sup> M |
|  |  | 8x10<sup>-8</sup> | 7x10<sup>-12</sup> M |
|  |  | 8x10<sup>-10</sup> | 7x10<sup>-14</sup> M |
| Carvone | CAR | 4x10<sup>-2</sup> | 1x10<sup>-7</sup> M |
| Ethyl tiglate | ETG | 3x10<sup>-2</sup> | 1x10<sup>-6</sup> M |
| Hexanal | HEX | 1.1x10<sup>-2</sup> | 1x10<sup>-6</sup> M |
| Benzal-dehyde | BZD | 8x10<sup>-2</sup> | 1x10<sup>-7</sup> M |

### Odor delivery

To deliver odor stimulus for both electrophysiological and behavioral experiments, to deliver odor stimulus, we used an eight-odor air dilution olfactometer (Fig.1A). Approximately 1 sec prior to odor delivery, a stream of Nitrogen was diverted through one of the odorant vials at a rate between 100 and 10 ml/min, and then merged into a clean air stream, flowing at a rate between 900 and 990 ml/min, thus providing 10 to 100 fold air dilution. Gas flows were controlled by mass flow controllers (Alicat MC series) with 0.5% accuracy. The odorized stream of 1000 ml/min was homogenized in a long thin capillary before reaching the final valve. Between stimuli, a steady stream of clean air with the same rate flowed to the odor port continuously, and the flow from the olfactometer was directed to an exhaust.

During stimulus delivery, a final valve (four-way Teflon valve, NResearch, SH360T042) switched the odor flow to the odor port, and diverted the clean airflow to the exhaust ([38], Suppl. Fig 1). Temporal odor concentration profile was checked by mini-PID (Aurora Scientific, model 200B). The concentration reached a steady state 95–210 ms (depending on a specific odor) after final valve opening. To minimize pressure shocks and provide temporally precise, reproducible, and fast odor delivery, we matched the flow impedances of the odor port and exhaust lines, and the flow rates from the olfactometer and clean air lines. A custom Phyton code monitored sniff pressure in real time and controlled the opening of the final valve at the onset of exhalation, so that the odor reached steady-state concentration before the next inhalation. At the end of the odor delivery (duration 1-4 s) the final valve was deactivated, and Nitrogen flow was diverted from the odor vial to an empty line. Inter-odor delivery interval was 7-14 s, during which clean air was flowing through all Teflon tubing.

### Sniff recording

To monitor the sniff signal, A sniffing cannula located in the nose port was connected to a pressure sensor through an 8-12 cm long polyethylene tube (801000, A-M Systems). The pressure was transduced with a pressure sensor (24PCEFJ6G, Honeywell) and preamplifier circuit. The signal from the preamplifier was recorded together with electrophysiological data on one of the data acquisition channels.

### Light stimulation

Light stimulation was produced via a 100 µm multimodal fiber coupled to a 473-nm diode laser (model FTEC2471-M75YY0, Blue Sky Research). The end of the fiber was cut flat and polished. The light stimulus power at the open end was measured by a power meter (Model, PM100D, Thorlabs), and calibrated to adjust the amplitude of the voltage pulses sent to the laser, to achieve a consistent power output across experiments.

### Electrophysiology

A bespoke array of PEDOT:PSS microelectrodes on parylene C was developed for this work using a previously reported fabrication process [29], with electrodes that had an area of 18×18 µm^2^. Neural signals were recorded using 64 channels digital headstages (Intan RHD-2000, Intan Technologies California, USA) and electrophysiology system [39] (Open ephys inc., Massachusetts, USA). Signals were recorded at 2 KHz frequency. The analog sniff pattern signal, and multiple triggers, such as a final valve opening and a beginning of the trail, were synchronously recorded with neural data all as 0-5V analog signals.

### Data analysis

#### Preprocessing

All data analysis was performed in Matlab (The MathWorks, Natick, MA). A 60Hz notch filter was applied to all raw signals to remove AC line voltage noise. All signals were low-pass filtered (<100 Hz, 4^th^ order Butterworth filter) and downsampled (10-fold, 200 Hz). Single electrode signals with peaks exceeding 2 mV in a period of 5 s were considered damaged and excluded from the study.

Odor presentation onset was defined as the first time point after the final valve opening when the sniff pressure signal crossed the baseline threshold, indicating inhalation onset. The raw trial signals were then segmented into individual trials, each starting 1 sec before inhalation onset, and ending 2.5 seconds after inhalation onset.

The common signal was calculated as the mean response across all electrode sites. Site-specific responses were obtained by subtracting this common signal from the responses at individual sites.

#### Dimensionality reduction

To reduce the dimensionality of the multi electrode signal, we performed principal component analysis (PCA). 5 PCs were used, explaining 87%±4.6% (s.e.m) of the variance.

#### Classification of odor identity

Decoding of odor identity was performed using a support vector machine (SVM) with a linear kernel. Feature vectors were built by concatenating different signals measured in a time window of 300ms after stimulus onset, discretized in 5ms bins. Signals included the common signal and/or site-specific signals, in their original form, or projected in the space of the first 5 PCs. For all results we report cross-validated classification performance (5-fold), with variance estimated via bootstrapping (across 100 repetitions).

#### Estimation of odor concentration

Estimation of odor concentration was performed using multivariate linear regression, with the same feature vectors as described above, with 5-fold cross-validation.

#### Calculation of detection accuracy: neural signals

Using the recording for PEA and MVT at five different concentrations (see Table 1), we trained linear kernel SVMs to discriminate odors from air, separately for each concentration, using both the common and site-specific signals as features. Performance was estimated using 5-fold cross validation. The same procedure was used for discriminating odors vs. air, in the presence of a background odor, at low/high concentration.

#### Calculation of detection accuracy

The same set of stimuli was used to measure behavioral detection thresholds (see [16] for detailed description). As in the original paper, behavioral performance was fit as:

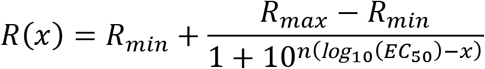

where *x* log_10_ of the concentration, *R_min_* and *R_max_* mark minimal and maximal responses, respectively, EC_50_ is the concentration at half maximal response, and *n* is the Hill slope. The parameters were estimated using nonlinear regression. The coefficients were estimated using iterative least square estimation. (Matlab nlinfit). P-values in fig. 4 were calculated using two-tailed t-test (Matlab)

## ACKNOWLEDGEMENTS

We thank Ezequiel Arneodo and Dion Khodagholy for a help at the initial stage of the project, David Godovich, Alexandra Dolzhina, and members of the Rinberg lab for technical help. The project was funded by DARPA BAA 15-35. PHV was supported by training grant R90DA043849 (NIH). The fabrication of the electrodes was supported by the European Union’s Horizon 2020 Research and Innovation Programme under grant agreement No. 732032 (BrainCom).

## AUTHOR CONTRIBUTIONS

E.S., T.B, and D.R. conceived and designed the experimental approaches. E.S. and P.H-V collected the electrophysiological data, A.D. collected behavioral data, P.H-V, E.S. and C.S. performed data analysis, T.B. generated transgenic mice, I.U., V.F.C. and G.M developed and provided electrodes, E.S., P.H-V, C.S., T.B, and D.R wrote the manuscript, D.R. supervised the project.

## SUPPLEMENTARY FIGURES

**Supplementary Figure 1.**
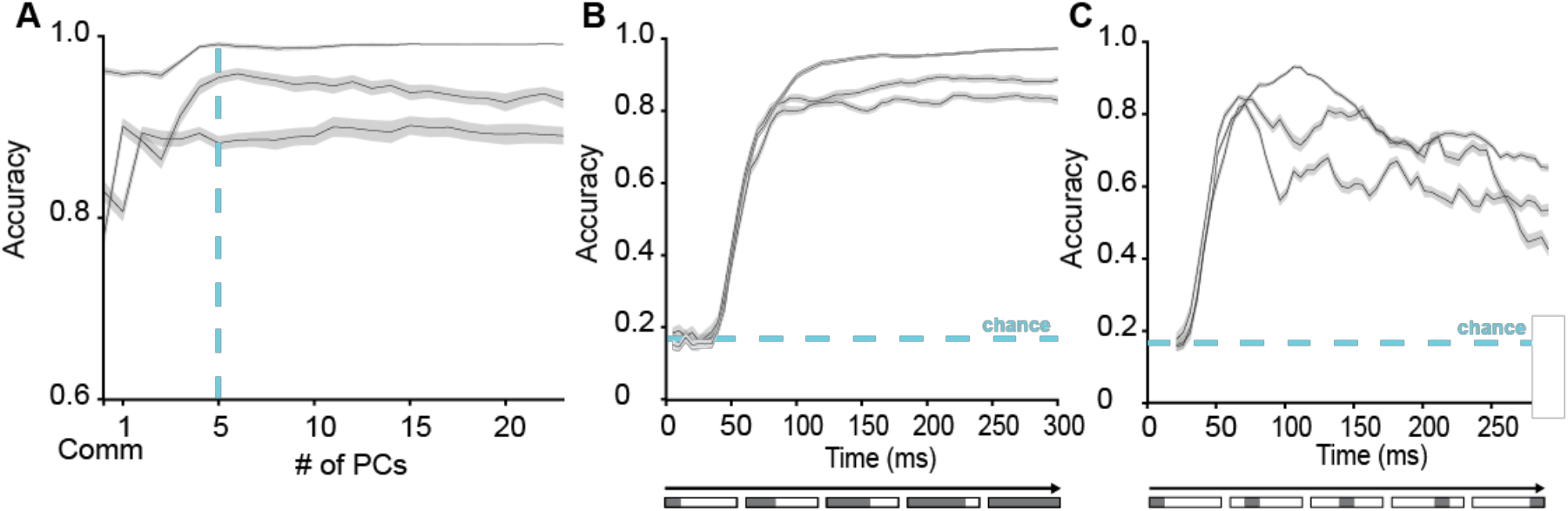
**A.** Average classification accuracy when varying the number of PC dimensions included in the feature vector, for n=3 representative mice. **B.** Classification accuracy as a function of window duration, for n=3 representative mice. **C**. Classification accuracy for a 30 ms sliding window, for n=3 representative mice. All shaded areas mark standard deviation estimated using bootstrapping (n=100 samples).

**Supplementary Figure 2.**
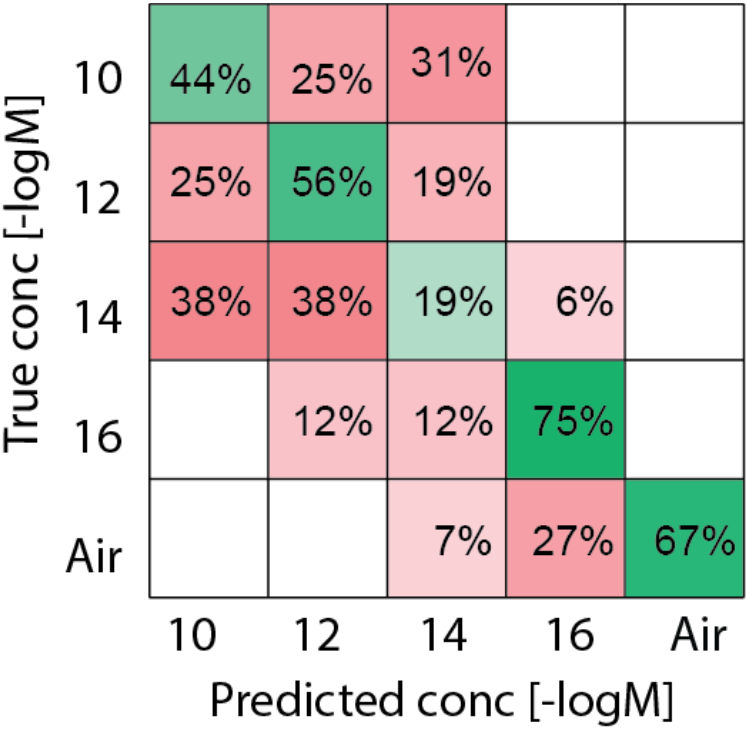
Confusion matrix for concentration classification (linear SVM, n = 79 trials).

